# Significantly different expression levels of microRNAs associated with vascular invasion in hepatocellular carcinoma and their prognostic significance after surgical resection

**DOI:** 10.1101/625095

**Authors:** Sung Kyu Song, Woon Yong Jung, Seung-Keun Park, Chul-Woon Chung, Yongkeun Park

## Abstract

**Background:** Although gross vascular invasion (VI) has prognostic significance in patients with hepatocellular carcinoma (HCC) who have undergone hepatic resection, few studies have investigated the relationship between gross VI and aberrant expression of microribonucleic acids (miRNAs and miRs). Thus, the objective of this study was to identify miRNAs selectively expressed in HCC with gross VI and investigate their prognostic significance.

**Materials and Methods:** Eligible two datasets (accession number: GSE20594 and GSE67140) were collected from the National Center for Biotechnology Information’s (NCBI) Gene Expression Omnibus (GEO) database to compare miRNAs expression between HCC with and without gross VI. Differentially expressed miRNAs were externally validated using expression data from The Cancer Genome Atlas (TCGA) database. Prognostic significance and predicted functions of selected miRNAs for HCC were also investigated.

**Results:** Thirty-five miRNAs were differentially expressed between HCC with and without gross VI in both datasets. Among them, four miRNAs were validated using TCGA database. miR-582 was upregulated to a greater extent while miR-99a, miR-100, and miR-148a were downregulated to a greater extent in patients with HCC and gross VI than in those with HCC but no VI. Receiver operating characteristic (ROC) curve analysis showed discriminatory power of these miRNAs in predicting gross VI. Multivariate survival analysis revealed that types of surgery, advanced tumor node metastasis (TNM) stage, and miR-100 underexpression were independently associated with tumor recurrence. It also revealed that types of surgery, advanced TNM stage, miR-100 underexpression, and miR-582 overexpression were independent risk factors for overall survival (OS) after hepatic resection for HCC. A text mining analysis revealed that these miRNAs were linked to multifaceted hallmarks of cancer, including “invasion and metastasis.”

**Conclusions:** miR-100 underexpression and miR-582 overexpression were associated with gross VI and poor survival of patients after hepatic resection for HCC.

## Introduction

Hepatocellular carcinoma (HCC) has received increasing attention because of its frequent diagnosis worldwide with a dismal prognosis [1]. Although various curative or palliative therapeutic modalities have been administered, long-term outcomes of patients with HCC have remained poor [2]. A significant contributor to poor outcomes is the tendency of HCC toward vascular invasion (VI) which reflects the tumor’s aggressiveness [3,4]. Therefore, altered or disrupted regulatory mechanisms contributing to VI in tumor cells a barrier to overcoming this cancer and so are candidate targets for a new therapeutic trial. These targets should be distinct genetic or pathologic features of HCC with VI definitely different from those of HCC without VI.

It is well known that unique gene expressions are highly related to tumor progression [5]. Recent studies have shown that aberrant epigenetic gene regulations also play a critical role in tumor progression [6]. Epigenetic alterations include deoxyribonucleic acid (DNA) methylation, histone modification, and ribonucleic acid (RNA) interference. Of these alterations, RNA interference can cause silencing of gene expression after introduction of sense–antisense RNA pairs [7]. Cells have a large number of noncoding RNA (ncRNA) molecules, many of which are capable of RNA interference. Recently, critical roles of small ncRNAs including microRNAs (miRNAs, miRs) and piwi-interacting RNAs (piRNAs) in a variety of human diseases, particularly cancers, have been well elucidated [8,9]. These RNAs have been shown to play an important role in tumorigenesis and/or tumor progression. Some of these RNAs can regulate epithelial–mesenchymal transition (EMT) directly or indirectly. Thus, they are involved in tumor invasion or metastasis [10]. Recent studies have identified clinically significant abnormal patterns of miRNA expression in HCC, with some miRNAs showing marked association with aggressive tumor phenotypes or poor survival [11,12].

Several studies have demonstrated correlations of VI and aberrant expression with specific miRNAs (down-regulation of miR-30a-3p and miR-34c-3p) [13,14]. However, these studies did not analyze all miRNA profiling with RNA sequencing technology or microarray analysis. They only focused on selected miRNAs in order to elucidate whether these miRNAs were distinctly expressed between HCC with VI and HCC without VI. Therefore, the objective of this study was to identify miRNAs that could act as important contributors to VI in HCC.

## Materials and methods

### Study data and screening of differentially expressed miRNAs

High-throughput miRNA expression data were obtained from the National Center for Biotechnology Information’s (NCBI) Gene Expression Omnibus (GEO) dataset [15] (http://www.ncbi.nlm.nih.gov/geo/) under accession numbers GSE20594 and GSE67140, including miRNA expression profiles of HCC tumor samples and information about VI status. Differentially expressed miRNAs between tumors on the basis of absence or presence of VI were detected with both Wilcoxon ranked-sum test with multiple test corrections (Benjamini–Hochberg-adjusted *p*-value < 0.05) and RankProd method [16]. Differentially expressed miRNAs between groups were validated and their prognostic values were additionally assessed using expression data obtained from The Cancer Genome Atlas (TCGA) database. Fig 1a shows a schematic flowchart of this study.

**Fig 1.**
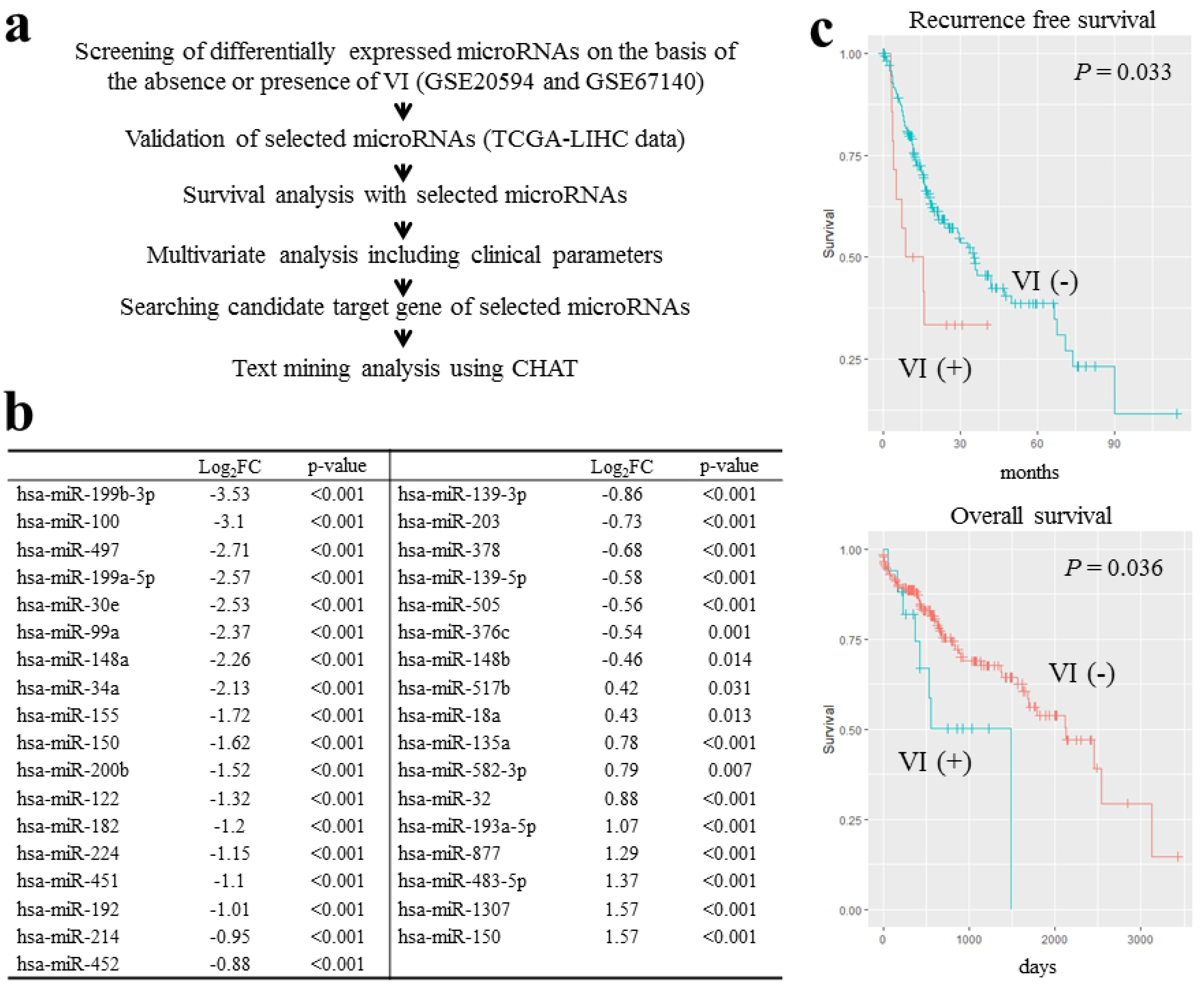
(a) Study flowchart, (b) Thirty-five differentially expressed miRNAs in gross VI group compared to negative VI group in both GSE20594 and GSE67140, (c) Comparison of RFS and OS of patients with gross VI [VI (+), group B] and without VI [VI (−), group A] in TCGA LIHC dataset. CHAT, Cancer Hallmarks Analytics Tool; TCGA LIHC, The Cancer Genome Atlas Liver Hepatocellular Carcinoma; VI, vascular invasion.

Clinical data for 377 patients with HCC were obtained from The Cancer Genome Atlas Liver Hepatocellular Carcinoma (TCGA LIHC) database (see https://tcga-data.nci.nih.gov and http://gdac.broadinstitute.org/) available to the public. Patients, their corresponding clinical data, and miRNA and messenger RNA (mRNA) expression profiles from the dataset were categorized into two groups on the basis of the absence of VI or presence of gross VI. Patients with missing gross VI values or with microVI (McVI) were included only for survival analysis according to the expression value of each candidate miRNA. Descriptive statistics were used to compare patient characteristics including sex, age, etiology, laboratory data, types of surgery, and pathologic data. A major hepatic resection was defined as the removal of three or more segments of the liver whereas a minor resection was defined as the removal of two segments or less.

MiRNA and mRNA expression data were generated using Illumina HiSeq 2000 sequencing platforms (Illumina Inc., San Diego, CA, USA). Raw read counts were used to analyze differentially expressed miRNAs and mRNAs. Processed RNA sequencing (RNA-Seq) data that were normalized according to reads per million (RPM) of miRNAs and mRNAs were also used in survival and correlation analyses. This study complied with publication guidelines provided by the TCGA (http://cancergenome.nih.gov/publications/publicationguidelines). All TCGA data are now available without restrictions on their use in publications or presentations. Differentially expressed sequence (DESeq2) package of the statistics software R version 3.3.1 was used to assess differentially expressed miRNAs on the basis of raw read counts [17]. Significantly different miRNA expression between groups was determined by false discovery rate (FDR) correction for multiple hypothesis testing using Benjamini–Hochberg-adjusted *p*-values with a threshold of FDR < 0.05. Receiver operating characteristic (ROC) curve analysis was performed to define optimal cutoff values of selected miRNAs on the basis of the absence or presence of gross VI.

### Survival analysis

Recurrence-free survival (RFS) and overall survival (OS) were estimated for both patient groups. We also calculated optimal cutoff values for continuous variables of clinical parameters for use in Kaplan–Meier survival analysis. They were estimated using ROC curve analysis. In addition, survival rates and curves were estimated by the Kaplan–Meier method and compared using the log-rank test. To calculate hazard ratios (HRs) and 95% confidence intervals (CIs) and identify independent factors associated with RFS and OS, multivariate analysis was performed using Cox regression proportional hazards model. All statistical analyses were performed using Epi, GGally, and Survival packages of R version 3.3.1 [18].

### Prediction of miRNA targets and text mining analysis

To identify target genes for the above candidate miRNAs, we obtained related genome data from NCBI GEO database which revealed changes in gene expression profile after overexpression of candidate miRNA mimic. Differentially expressed genes between control and candidate miRNA mimic-treated samples were analyzed using GEO2R (http://www.ncbi.nlm.nih.gov/geo/geo2r/). |log_2_FC| > 1 and adjusted *p*-value < 0.05 were used as cut-off criteria to define statistically significant difference. Selected genes were compared to a validated target module of MiRWalk2.0 database [19]. Only common genes were chosen for further analysis. We also performed Pearson’s correlation analysis to calculate correlations between the expression of candidate miRNAs and that of predicted target genes with all available samples in the TCGA LIHC database, filtering out genes with meaningful negative correlations (*r* < −0.3 and FDR < 0.05). Cancer Hallmarks Analytics Tool (CHAT) was used to figure out the association between selective miRNAs and documented evidence of these in hallmarks of cancer [20].

## Results

### miRNA differential expression analysis

As described in the detailed flowchart shown in Fig 1a, in this study, we obtained information about expression levels of miRNAs from NCBI GEO database [15]. To screen miRNAs that were significantly more or less abundant in HCC with VI compared to those in HCC without VI, we performed differential expression analysis by comparing the expression level of each miRNA from two microarray datasets (accession number: GSE20594 and GSE67140). These datasets included 31 samples of HCC without gross VI vs. 46 samples of HCC with gross VI (accession number: GSE20594) and 91 samples of HCC without gross VI vs. 81 samples of HCC with gross VI (accession number GSE67140). We then identified differentially expressed miRNAs based on the Wilcoxon ranked-sum test and the RankProd method [16]. A total of 49 miRNAs in GSE20594 were upregulated in HCC with VI compared to those in HCC without VI. In addition, 185 miRNAs in GSE67140 showed increased expression levels in HCC with VI compared to those in HCC without VI. Ten miRNAs were upregulated in these two microarray data sets. Forty-one miRNAs (GSE20594) and 184 miRNAs (GSE67140) were downregulated in HCC with VI compared to HCC without VI while 25 miRNAs were downregulated in both data sets. These 35 miRNAs were predicted to be candidate biomarkers for VI of HCC (Fig 1b). They were validated using expression data obtained from TCGA database.

### Data collection and overall evaluation of TCGA LIHC

We found no miRNA expression data for 5 of 377 patients with HCC in the TCGA LIHC database. The remaining 372 patients were divided into three groups by gross VI status as follows: group A (*n* = 206), patients with HCC without gross VI; group B (*n* = 17), patients with HCC and gross VI; and group C (*n* = 149), patients with HCC with McVI or patients for whom no gross VI status was available. Groups A and B were included in the study to find miRNAs expressed differentially between HCC with and without VI. Group C was included in survival analysis according to expression value of each candidate miRNA.

Table 1 summarizes clinical characteristics of groups A, B, and C. Group B had slightly higher preoperative platelet counts without statistical significance (*P* = 0.058). However, the percentage of patients who had undergone major hepatic resection was significantly lower (*P* < 0.001). Besides these differences, these three groups were comparable. The overall median follow-up period was 16 months (range, 0–122 months). Survival analysis showed significantly worse RFS (1-year RFS rates: group A, 75.7%; group B, 50.0%; *P* = 0.033) and OS (2-year OS rates: group A, 75.4%; group B, 50.3%; *P* = 0.036) for patients with HCC and gross VI (Fig 1c).

**Table 1.** Comparison of clinicopathological data of hepatocellular carcinoma patients with no vascular invasion or with gross vascular invasion

|  | Group A<br>(N=206) | Group B<br>(N=17) | Group C<br>(N=149) | P value |
| --- | --- | --- | --- | --- |
| Sex |  |  |  | 0.211 |
| Male | 136 (66.0%) | 9 (52.9%) | 107 (71.8%) |  |
| Female | 70 (34.0%) | 8 (47.1%) | 42 (28.2%) |  |
| Age | 59.9 ± 13.5 | 60.6 ± 14.7 | 58.6 ± 13.3 | 0.405 |
| Viral hepatitis |  |  |  | 0.176 |
| No | 81 (39.9%) | 10 (62.5%) | 65 (44.8%) |  |
| Yes | 122 (60.1%) | 6 (37.5%) | 80 (55.2%) |  |
| History of other malignancy |  |  |  | 0.417 |
| No | 185 (89.8%) | 14 (82.4%) | 137 (91.9%) |  |
| Yes | 21 (10.2%) | 3 (17.6%) | 12 (8.1%) |  |
| Platelet count (×1000/μL) | 206 ± 91 | 271 ± 85 | 228 ± 113 | 0.058 |
| Serum creatinine |  |  |  | 0.591 |
| W.N.L. | 173 (94.5%) | 14 (100.0%) | 97 (96.0%) |  |
| B.N.L. | 10 ( 5.5%) | 0 ( 0.0%) | 4 ( 4.0%) |  |
| Serum albumin |  |  |  | 0.635 |
| W.N.L. | 138 (75.8%) | 13 (86.7%) | 78 (76.5%) |  |
| B.N.L. | 44 (24.2%) | 2 (13.3%) | 24 (23.5%) |  |
| Serum total bilirubin |  |  |  | 0.456 |
| W.N.L. | 156 (84.8%) | 16 (94.1%) | 84 (82.4%) |  |
| B.N.L. | 28 (15.2%) | 1 ( 5.9%) | 18 (17.6%) |  |
| Child-Turcotte-Pugh classification |  |  |  | 0.416 |
| A | 144 (90.0%) | 11 (84.6%) | 66 (94.3%) |  |
| B or C | 16 (10.0%) | 2 (15.4%) | 4 ( 5.7%) |  |
| Alpha-fetoprotein (ng/mL) | 15292.7 ±<br>157568.5 | 16856.2 ±<br>34681.1 | 10958.2 ±<br>42145.9 | 0.789 |
| Types of hepatic resection |  |  |  | < 0.001 |
| Major | 133 (64.6%) | 9 (52.9%) | 57 (39.0%) |  |
| Minor | 73 (35.4%) | 8 (47.1%) | 89 (61.0%) |  |
| Histologic grading by Edmondson and Steiner's classification |  |  |  | 0.384 |
| 1~2 | 131 (63.9%) | 8 (47.1%) | 92 (63.0%) |  |
| 3~4 | 74 (36.1%) | 9 (52.9%) | 54 (37.0%) |  |
| Residual tumor |  |  |  | 0.249 |
| Negative | 181 (96.3%) | 15 (88.2%) | 129 (93.5%) |  |
| Positive | 7 ( 3.7%) | 2 (11.8%) | 9 ( 6.5%) |  |
| Ishak fibrosis staging system |  |  |  | 0.295 |
| 0 | 52 (36.9%) | 5 (55.6%) | 17 (26.2%) |  |
| 1~4 | 36 (25.5%) | 2 (22.2%) | 23 (35.4%) |  |
| 5~6 | 53 (37.6%) | 2 (22.2%) | 25 (38.5%) |  |
| Grade of liver inflammation |  |  |  | 0.139 |
| 0 | 64 (46.4%) | 9 (81.8%) | 44 (50.6%) |  |
| 1~2 | 65 (47.1%) | 2 (18.2%) | 34 (39.1%) |  |
| 3~4 | 9 ( 6.5%) | 0 ( 0.0%) | 9 (10.3%) |  |

Notes: Group A, no vascular invasion; Group B, gross vascular invasion; Group C, microvascular invasion or no available information regarding vascular invasion. Mismatches of patients numbers across the variables occurs due to some missing values. B.N.L.: Beyond normal range; W.N.L.: With normal range.

### MiRNAs differentially expressed on the basis of the absence or presence of gross VI in TCGA database

We obtained information about expression levels of miRNAs from TCGA expression data. A total of 26 miRNAs were differentially expressed between group A and group B, and 4 miRNAs were on the above list of candidate miRNAs from two GEO datasets (Fig 1b). miR-99a, miR-100, and miR-148a were downregulated to a greater extent in group B (log_2_ FC = −1.12 with FDR = 0.009, log_2_ FC = −1.45 with FDR < 0.001, and log_2_ FC = −0.91 with FDR < 0.011) while miR-582 was upregulated to a greater extent in group B (log_2_ FC = 1.09 with FDR = 0.015) (Fig 2a).

**Fig 2.**
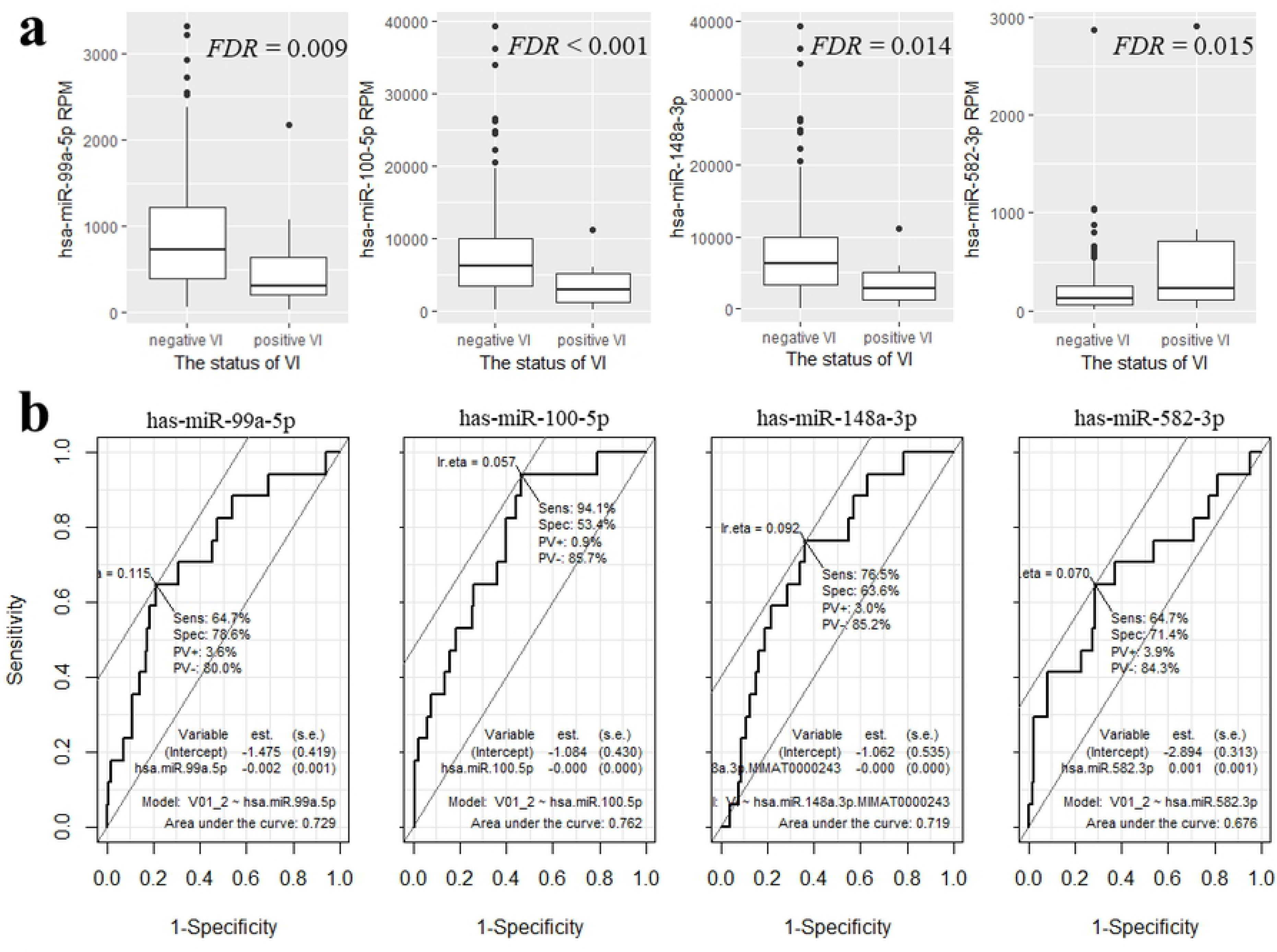
(a) Expression difference of four miRNAs according to vascular invasion status, and (b) ROC curves of these miRNAs distinguishing HCC tissues on the basis of the absence or presence of gross VI (miR-99a-5p, miR-100-5p, miR-148-3p and miR-582-3p, respectively). FDR, false discovery rate.

Next, we assessed the discriminatory power of the remaining three miRNAs in predicting gross VI. To evaluate the predictive value, we used ROC curve to analyze sensitivity and specificity. As shown in Fig 2b, the ROC curve of miR-99a-5p showed an area under the curve (AUC) of 72.9% (sensitivity, 65.7%; specificity, 78.6%), similar to the AUC of miR-100-5p at 76.2% (sensitivity, 94.1%; specificity, 53.4%) and the AUC of miR-148a-3p at 71.9% (sensitivity, 76.5%; specificity, 63.6%). miR-582-3p yielded a ROC curve AUC of 67.6% (sensitivity, 64.7%; specificity, 71.4%). HCC with gross VI showed at least one aberrant expression pattern in these four miRNAs.

### Survival analysis

ROC-curve-determined optimal cutoff value of miRNA expression in study participants (n = 372) was used to classify patients into under- and overexpression groups. Cutoff values for miR-99a-5p, miR-100-5p, miR-148a-3p, and miR-582-3p expression were 350.9, 5964.4, 101299.6, and 219.7 RPM, respectively. Kaplan–Meier analysis indicated that underexpression of miR-100-5p and miR-148a-3p and overexpression of miR-582-3p were significantly associated with poor RFS (log-rank test: *P* < 0.001, *P* = 0.024, and *P* = 0.009, respectively) and worse OS (log-rank test: *P* < 0.001, *P* = 0.002, and *P* < 0.001, respectively) (Table 2). However, miR-99a-5p underexpression showed a significant difference in OS outcome (log-rank test: *P* < 0.001), but not in RFS outcome.

**Table 2.** Univariate analysis of factors predictive of recurrence-free survival and overall survival

| Factors | No. of patients | MDFST<sup>1</sup> (95% CI) | *P* value | Factors | No. of patients | MOST<sup>2</sup> (95% CI) | *P* value |
| --- | --- | --- | --- | --- | --- | --- | --- |
| **Sex** |  |  | 0.617 | **Sex** |  |  | 0.244 |
| Male | 222 | 21.2 (16.5–33.0) |  | Male | 250 | 1791 (1423-NA) |  |
| Female | 100 | 19.2 (14.8–35.8) |  | Female | 120 | 1450 (899-2456) |  |
| **Age (years)** |  |  | 0.659 | **Age (years)** |  |  | 0.127 |
| <43 | 33 | 15.7 (8.6–NA) |  | <69 | 275 | 2456 (1622-NA) |  |
| ≥43 | 289 | 21.2 (18.3–29.3) |  | ≥69 | 95 | 1386 (887-1836) |  |
| **Hepatitis B or C infection status** |  |  | 0.510 | **Hepatitis B or C infection status** |  |  | 0.018 |
| Negative | 126 | 19.2 (15.7–33.0) |  | Negative | 154 | 1135 (837-1694) |  |
| Positive | 188 | 21.6 (16.5–35.8) |  | Positive | 208 | 2456 (1624-NA) |  |
| **Prothrombin time (sec.)** |  |  | 0.474 | **Prothrombin time (sec.)** |  |  | 0.474 |
| With normal range | 220 | 23.0 (19.2-35.8) |  | With normal range | 248 | 2116 (1560-NA) |  |
| Beyond normal range | 40 | 29.3 (18.3-NA) |  | Beyond normal range | 44 | NA(NA-NA) |  |
| **Serum creatinine (mg/dL)** |  |  | 0.564 | **Serum creatinine (mg/dL)** |  |  | 0.564 |
| With normal range | 251 | 23.0 (19.2–33.9) |  | With normal range | 283 | 2131 (1622-NA) |  |
| Beyond normal range | 10 | 45.8 (11.8–NA) |  | Beyond normal range | 14 | 785 (419-NA) |  |
| **Serum albumin (g/dL)** |  |  | 0.622 | **Serum albumin (g/dL)** |  |  | 0.622 |
| With normal range | 206 | 21.6 (18.3-35.6) |  | With normal range | 228 | 2116 (1560-NA) |  |
| Beyond normal range | 58 | 25.5 (15.9-71.1) |  | Beyond normal range | 70 | NA (837-NA) |  |
| **Serum total bilirubin (mg/dL)** |  |  | 0.145 | **Serum total bilirubin (mg/dL)** |  |  | 0.145 |
| With normal range | 230 | 24.8 (19.4-36.7) |  | With normal range | 256 | 2131 (1624-3258) |  |
| Beyond normal range | 37 | 25.5 (14.8-NA) |  | Beyond normal range | 46 | 1622 (785-NA) |  |
| **Child–Turcotte–Pugh classification** |  |  | 0.153 | **Child–Turcotte–Pugh classification** |  |  | 0.153 |
| A | 196 | 23.6 (18.6–36.7) |  | A | 220 | 2542 (1694-NA) |  |
| B or C | 19 | 23.9 (8.6–NA) |  | B or C | 22 | 887 (601-NA) |  |
| **Alpha-fetoprotein (ng/mL)** |  |  | 0.085 | **Alpha-fetoprotein (ng/mL)** |  |  | 0.085 |
| <103,900 | 240 | 24.8 (19.3–35.8) |  | <85,150 | 270 | 2456 (1624-NA) |  |
| ≥103,900 | 7 | NA (7.9–NA) |  | ≥85,150 | 10 | NA (NA-NA) |  |
| **Extent of resection** |  |  | <0.001 | **Extent of resection** |  |  | 0.024 |
| Major | 169 | 35.6 (23.0–53.5) |  | Major | 198 | 2116 (1624-NA) |  |
| Minor | 152 | 13.6 (9.89–19.2) |  | Minor | 169 | 1147 (848-NA) |  |
| **Residual tumor** |  |  | 0.116 | **Residual tumor** |  |  | 0.168 |
| No | 288 | 21.6 (18.2–33.0) |  | No | 324 | 1791 (1450-2542) |  |
| Yes | 15 | 15.6 (10.8–NA) |  | Yes | 18 | 1135 (837-NA) |  |
| **Histologic grading by Edmondson and Steiner's classification** |  |  | 0.421 | **Histologic grading by Edmondson and Steiner's classification** |  |  | 0.654 |
| I~II | 200 | 21.6 (18.4–30.0) |  | I~II | 230 | 1791 (1386-2542) |  |
| III~IV | 118 | 15.7 (11.7–42.0) |  | III~IV | 136 | 1622 (1149-NA) |  |
| **Ishak fibrosis staging system** |  |  | 0.472 | **Ishak fibrosis staging system** |  |  | 0.472 |
| 0 | 60 | 23.9 (15.7–NA) |  | 0 | 74 | 2131 (931-NA) |  |
| 1–4 | 57 | 24.8 (16.5–36.7) |  | 1–4 | 61 | 1791 (1372-NA) |  |
| 5–6 | 73 | 18.3 (12.9–47.0) |  | 5–6 | 79 | NA (1685-NA) |  |
| **Grade of liver inflammation** |  |  | 0.369 | **Grade of liver inflammation** |  |  | 0.369 |
| 0 | 104 | 23.9 (18.6–40.4) |  | 0 | 117 | 2131 (1624-NA) |  |
| 1–2 | 92 | 21.6 (17.6–47.0) |  | 1–2 | 101 | NA (1372-NA) |  |
| 3–4 | 15 | 18.3 (9.7–NA) |  | 3–4 | 17 | NA (359-NA) |  |
| **American Joint Committee on Cancer TNM stage** |  |  | <0.001 | **American Joint Committee on Cancer TNM stage** |  |  | <0.001 |
| I~II | 225 | 30.0 (23.6–42.0) |  | I~II | 257 | 2456 (1694-NA) |  |
| III~IV | 76 | 9.6 (8.3–14.2) |  | III~IV | 89 | 660 (445-1791) |  |
| **miR-99a** |  |  | 0.144 | **miR-99a** |  |  | <0.001 |
| Under-expression | 223 | 18.4 (14.2-28.9) |  | Under-expression | 123 | 837 (556-NA) |  |
| Over-expression | 99 | 23.6 (20.9-71.1) |  | Over-expression | 247 | 2116 (1560-NA) |  |
| **miR-100** |  |  | <0.001 | **miR-100** |  |  | <0.001 |
| Under-expression | 71 | 9.9 (7.2-15.7) |  | Under-expression | 87 | 639 (421-1490) |  |
| Over-expression | 251 | 27.2 (20.9-36.7) |  | Over-expression | 283 | 2131 (1624-3258) |  |
| **miR-148a** |  |  | 0.024 | **miR-148a** |  |  | 0.002 |
| Under-expression | 143 | 24.8 (21.0-47.7) |  | Under-expression | 185 | 2542 (1685-NA) |  |
| Over-expression | 179 | 16.4 (13.1-25.3) |  | Over-expression | 185 | 1372 (931-1791) |  |
| **miR-582** |  |  | 0.009 | **miR-582** |  |  | <0.001 |
| Under-expression | 143 | 28.9 (21.2–47.0) |  | Under-expression | 312 | 1791 (1490-3258) |  |
| Over-expression | 179 | 15.7 (12.6–23.6) |  | Over-expression | 58 | 757 (415-NA) |  |

**Notes:** <sup>1</sup>Median disease-free survival time (month), <sup>2</sup>Median overall survival time (day). Mismatches of patients numbers across the variables occurs due to some missing values.
**Abbreviations:** TNM (Tumor Node Metastasis), CI (confidence interval), NA (not available).

### Multivariate analysis

In this study, to identify possible predictors of tumor recurrence and poor survival of patients with HCC after hepatic resection, we analyzed other associated clinicopathological factors by univariate and multivariate methods (Table 2). Univariate analysis showed significant associations of tumor recurrence with types of surgery (*P* < 0.001) and advanced TNM stage (*P* < 0.001). Multivariate logistic regression was performed for each predictor of recurrence, including gross VI, miR-100-5p underexpression, and miR-582-3p overexpression. We found that types of surgery (HR = 1.676; 95% CI: 1.205–2.332; *P* = 0.002), advanced TNM stage (HR = 1.795; 95% CI: 1.266–2.547; *P* = 0.001), and miR-100-5p underexpression (HR = 1.511; 95% CI: 1.041–2.194; *P* = 0.029) remained significant independent risk factors for tumor recurrence (Fig 3). Types of surgery, concomitant viral hepatitis, and advanced TNM stage were associated with poorer OS in univariate Kaplan–Meier survival estimation with log-rank comparisons (*P* = 0.024, *P* = 0.019, and *P* < 0.001, respectively). The multivariate proportional hazards model identified that types of surgery (HR = 1.844; 95% CI: 1.110–3.065; *P* = 0.018), advanced TNM stage (HR = 2.401; 95% CI: 1.433–4.024; *P* < 0.001), miR-100-5p underexpression (HR = 2.652; 95% CI: 1.519–4.629; *P* < 0.001), and miR-582-3p overexpression (HR = 2.016; 95% CI: 1.109–3.663; *P* = 0.021) were independent risk factors for OS after hepatic resection in HCC (Fig 3).

**Fig 3.**
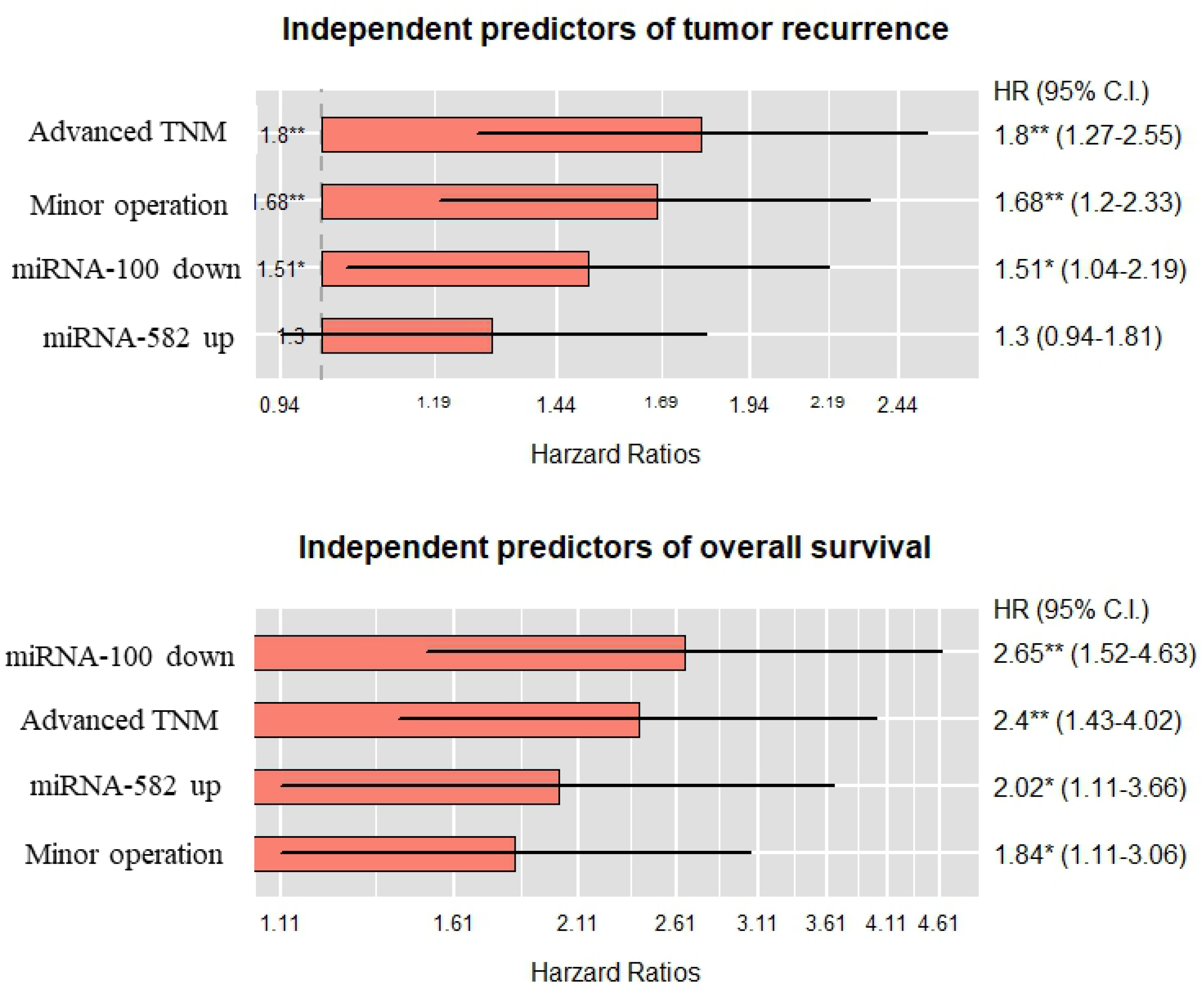
Summary of statistically significant factors on tumor recurrence and overall survival using Cox regression proportional hazards model. TNM, American Joint Committee on Cancer Tumor Node Metastasis; HR, hazard ratio; CI, confidence interval.

### Prediction of miRNA targets and text mining analysis

To identify target genes for miR-100-5p, we obtained two published microarray data from the NCBI website which revealed changes of gene expression profile after overexpression of miR-100-5p mimic (GEO accession numbers: GSE16571 and GSE20668). Each dataset included three biological replicates for both control and miRNA mimics treated cells (a total of six samples: GSE16571) and two replicates (a total of 4 samples: GSE20668). A total of 1181 genes were significantly down-regulated in miR-100-5p mimic treated samples (log_2_ FC < −1 and adjusted *p*-value < 0.05) in GSE16571 dataset. However, there was no gene expression change fulfilling the cut-off criteria in GSE20668 dataset. We also found a list of 243 predicted target genes of miR-100-5p using miRNA– mRNA pairs experimentally validated in the MiRWalk2.0 database [19]. The overlap of both lists comprised 21 genes (Fig 4a). Next, we performed Pearson’s correlation analysis to calculate correlations between the expression of miR-100-5p and of the above 21 genes with all available 424 samples (both mRNA and miRNA) from the TCGA LIHC database. Among 21 genes, expression values of *ACSL3* and *CTDSPL* had significant negative correlations with miR-100-5p (Pearson’s *r* = – 0.310 and –0.330, respectively, FDR < 0.001, Fig 4b). There was no available dataset for finding targets of either miR-148a or miR-582-3p from NCBI GEO database.

**Fig 4.**
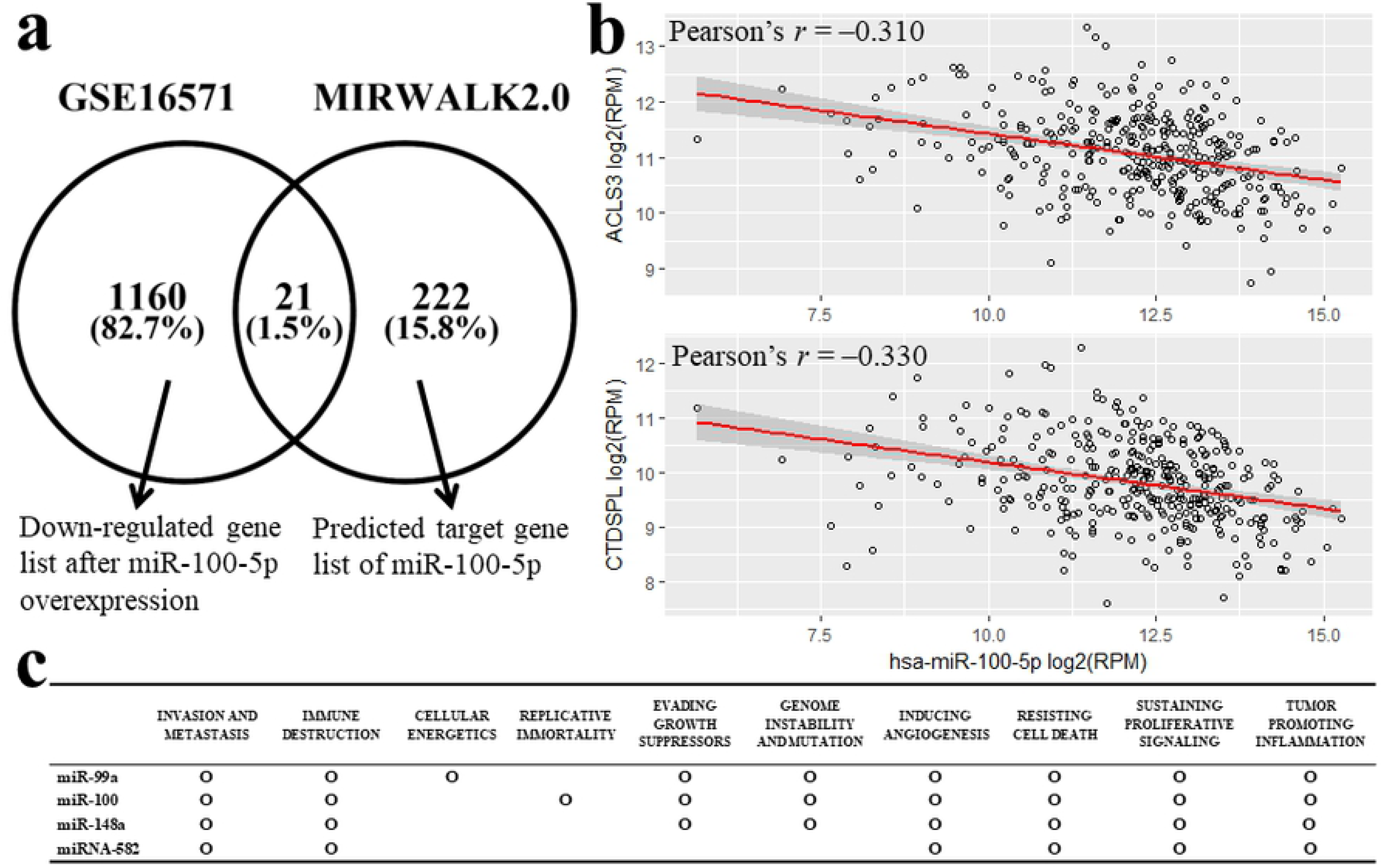
(a) Venn diagram demonstrating the overlap between differential mRNAs screened by GSE16571 dataset and miR-100-5p targets from MiRWalk2.0 database, and (b) Scatter plot showing significantly negative correlation between miR-100-5p and mRNA expression levels of *ACLS3* and *CTDSPL*. (c) Summary of CHAT analysis for miR-99a, miR-100, miR-148a, and miR-582. RPM, reads per million.

A text mining analysis was conducted to determine which hallmarks of cancer were affected when miR-99a/-100/-148a/-582 were dysregulated. All miRNAs were associated with at least six hallmarks of cancer. Especially, invasion and metastasis, immune destruction, inducing angiogenesis, resisting cell death, sustaining proliferative signaling, and tumor promoting inflammation were affected by all three miRNAs (Fig 4c).

## Discussion

In this study, we used publicly available data from TCGA repositories to find specific miRNAs associated with gross VI in HCC. We found that down-regulation of miR-99a/-100/-148a and up-regulation of miR-582 were commonly associated with gross VI in three publicly open datasets (TCGA and GEO, accession numbers: GSE20594 and GSE67140). Specifically, aberrant expressions of two miRNAs (miR-100 and miR-582) were independent risk factors for HCC recurrence and/or survival after hepatic resection for HCC.

Tumor biology research has focused on miRNAs. Many studies have demonstrated the involvement of miRNAs in cancer metastasis [21]. Compared with parenchyma tumors of HCC, one study has shown increased expression of miR-135a in a portal vein tumor thrombus [22]. However, it is difficult to find studies comparing expression values of all miRNAs on the basis of the absence or presence of gross VI. To the best of our knowledge, no previous study has investigated the association between aberrant expression of these four miRNAs (miR-99a, miR-100, miR-148a, and miR-582) and gross VI in HCC tissue samples.

Gross VI is a well-known factor that contributes to tumor recurrence and poor prognosis. A recent study has reported that its prevalence in newly detected HCC is 35% [23]. Notably, gross VI is found to be a major prognostic factor following hepatic resection for HCC [24]. Unfortunately, the optimal treatment strategy for HCC with gross VI remains inconclusive. The current Barcelona clinic liver cancer staging system recommends systemic therapy with sorafenib at this stage. However, outcome following this treatment is not promising [25]. Under careful selection based on good hepatic reserve and favorable preoperative predictors, hepatic resection has been justified in patients with HCC and gross VI [26]. Although hepatic resection has clinical significance, there is limited information about the underlying mechanism for gross VI. One possible inference is that gross VI merely occurs coincidentally, with high arterial pressure in the tumor acting as a driving force of cancer cells to neighboring portal or venous vessels. However, recent genomic data analyses have demonstrated that unique genes and ncRNAs play an important role in metastasis [27].

miRNA is one of endogenously expressed small ncRNAs (~22 nucleotides in length) with capacity as posttranscriptional regulators through degradation of their target mRNA or by inhibiting translation [28]. miRNA has been known to have a regulatory role in tumor progression such as migration, angiogenesis, and invasion as a powerful regulator of critical biological processes, including cell cycle, metabolism, development, and cell differentiation [13,29]. miR-100 has been shown to be a prognostic factor for different types of cancer [30]. One study has concluded that miR-100 down-regulation is correlated with progressive pathological features and poor prognosis in patients with HCC [31]. However, the exact role of miR-100 in preventing cancer progression has not been fully elucidated yet, although its importance has been recognized. Recently, miR-100 has been shown to be involved in suppressing migration and invasion of nasopharyngeal carcinoma cells by targeting insulin-like growth factor 1 receptor [32]. In our study, mRNA expressions of *ACSL3* and *CTDSPL* showed significant negative correlations with miR-100, consistent with results of an earlier study showing that the expression of *CTDSPL* (also known as *RBSP3*) protein had the most inverse correlation with miR-100 in clinical samples of acute myeloid leukemia [33]. Altered fatty acid metabolism in cancer has been increasingly recognized. Long-chain acyl-CoA synthetases (ACSLs) with responsibility for activation of long-chain fatty acids are commonly deregulated in cancer. Among ACSL family, ACSL3 was found to be overexpressed in different types of cancer [34]. However, the prognostic role of ACSLs remains unclear.

In the present study, miR-582 up-regulation was associated with gross VI. It showed a statistically significant difference in survival. A previous study has reported that miR-582 can promote cancer stem cell traits of non-small-cell lung cancer while miR-582 inhibition can potently inhibit tumor progression [35]. Therefore, these two miRNAs warrant attention as potential therapeutic targets for cancer treatment.

Recently, a similarly designed study using the TCGA database for investigating gross VI– related miRNAs expression profiles has been published [36]. Results of that study concluded that 16 miRNA-based classifiers could effectively identify gross VI. They also play a role as prognostic factors. However, miRNAs selected in that study were completely different from those selected in our study. In the TCGA LIHC database, we can recognize two types of VI from the pathologic data of every patient, McVI and gross VI. There is no mention about such information in previous studies. This is why different miRNAs are sorted. Because there is little evidence that McVI can be considered the first step of metastatic dissemination via the vascular route and McVI will advance to gross VI, we thought it was not justified that McVI and gross VI seemed to be the same pathobiological feature. Also, the prognostic impact of McVI remains inconclusive [37,38]. Therefore, patients with HCC with McVI were excluded from analysis in our study.

Over the past two decades, there has been a dramatic increase in genomic databases for diseases, including those for cancer. The TCGA provides a large volume of publicly available cancer genomic data. Researchers interested in molecular biology of cancer can access this valuable source of data. In the present study, we used epigenetic profiles of each HCC tumor sample with clinical parameters, enabling the establishment of new knowledge about the role of miRNAs in HCC with gross VI. In the near future, it is likely that newly discovered molecular targets based on the TCGA database will be applied to clinical cancer practice, including for early detection, treatment, and even prevention of cancers. However, effective translation of cancer genomics or proteomics to clinical practice requires progress in analytics, which in turn, requires close cooperation between bioinformaticians, mathematicians, and oncologists.

A limitation of our study was that it was based on secondary data analysis. Therefore, there was a lack of information about important perioperative data such as liver enzyme profiles, antiviral drug use, and postoperative progression of underlying liver disease. These are believed to influence tumor recurrence and *de novo* malignancy. Additionally, data on tumor size and the number of tumors in each patient are omitted from the TCGA database, although these two parameters are considered important prognostic factors. Unlike miR-100, no target candidate gene of miR-582 could be identified despite its prognostic significance. It is known that miRNAs can inhibit target genes via degradation of their target mRNAs or by inhibition of translation. If a specific miRNA functions through suppressing translation but not through degrading mRNA, it is difficult to find a target just from miRNA and mRNA expression profiles and their correlation analysis. In the TCGA database, protein expression data are limited compared with miRNA and mRNA data. Therefore, experimental verification is required in the future.

## Conclusion

This study showed that miR-100-5p was underexpressed while miR582-3p was overexpressed to a great extent in HCC tissues with gross VI than that in HCC tissues without gross VI. In addition, their aberrant expressions were significantly associated with poor survival of patients after hepatic resection for HCC.

